# Technical artifact drives apparent deviation from Hardy-Weinberg equilibrium at CCR5-Δ32 and other variants in gnomAD

**DOI:** 10.1101/784157

**Authors:** Konrad J. Karczewski, Laura D. Gauthier, Mark J. Daly

## Abstract

Following an earlier report suggesting increased mortality due to homozygosity at the CCR5-Δ32 allele^1^, Wei and Nielsen recently suggested a deviation from Hardy-Weinberg Equilibrium (HWE) observed in public variant databases as additional supporting evidence for this hypothesis^2^. Here, we present a re-analysis of the primary data underlying this variant database and identify a pervasive genotyping artifact, especially present at long insertion and deletion polymorphisms. Specifically, very low levels of contamination can affect the variant calling likelihood models, leading to the misidentification of homozygous individuals as heterozygous, and thereby creating an apparent depletion of homozygous calls, which is especially prominent at large insertions and deletions. The deviation from HWE observed at CCR5-Δ32 is a consequence of this specific genotyping error mode rather than a signature of selective pressure at this locus.

---

In a recent report, Wei and Nielsen^2^ use the Genome Aggregation Database (gnomAD) to show a depletion of homozygous individuals at the CCR5-Δ32 locus and interpret this deviation as evidence of negative selection against this allele. The authors use a control set of variants to show that this variant is nominally significantly depleted for homozygotes; however, systematic errors or missingness patterns that are non-uniformly distributed across genotype categories would show a similar signal. Indeed, deviation from Hardy-Weinberg Equilibrium (HWE) is often used as a part of the quality control process of large sequencing datasets and thus careful interpretation of these signals is warranted. Here, we identify and characterize an error mode in the gnomAD dataset (and many other datasets joint-called by GATK) that leads to this apparent deviation.

The CCR5-Δ32 frameshift variant is directly genotyped in the gnomAD exome and genome datasets^3^, with a call rate of 99.8%. In five subpopulations, the frequency of the variant is at least 10% (Finnish, Swedish, Estonian, North-Western European, Other Non-Finnish European) and the original report considers each of these as well as the combination of these populations. All but the Finnish cohort show a slight depletion of homozygous individuals in the frequencies described in the release dataset, with the Swedish cohort and the full dataset reaching nominal significance by at least one statistical test. The authors of the original report acknowledge that systematic under-calling of homozygous minor allele individuals can cause such a deficiency; however, it can be difficult to accurately characterize the underlying factors that lead to these artifacts.

A close analysis of the variant metadata in gnomAD reveals that 59 of the called heterozygotes in these populations have allele balance greater than 0.9 (i.e. greater than 90% of the reads support the alternate allele), suggesting that these individuals may be misclassified homozygotes for the CCR5-Δ32 allele (i.e. undercalled genotyping errors). This variant information is provided in the VCF as well as the browser, though not overtly flagged and not broken down by population. Recharacterizing these individuals as homozygous attenuates the signal in all populations, resulting in no overall excess of homozygotes, rather than a 5.4% depletion as previously reported (Table 1).

**Table 1.** Recomputation of Hardy-Weinberg Equilibrium statistics. After reclassifying variants with greater than 0.9 allele balance as homozygotes, there is a slight global excess rather than depletion of homozygotes. Empirical p-value is generated as described previously^2^ as “P1” (the proportion of variants in a control set that have a more extreme B_i_ value than B_Δ32_).

| Population | Reported<br>homozygotes | Recomputed<br>homozygotes | Reported<br>$B_{\Delta 32}$ | Recomputed<br>$B_{\Delta 32}$ | p-value |
| --- | --- | --- | --- | --- | --- |
| FIN | 228 | 238 | 1.012 | 1.050 | 0.731 |
| SWE | 167 | 180 | 0.871 | 0.931 | 0.216 |
| EST | 25 | 33 | 0.782 | 1.002 | 0.507 |
| NWE | 302 | 323 | 0.939 | 0.997 | 0.438 |
| ONF | 171 | 178 | 0.947 | 0.982 | 0.312 |
| All | 893 | 952 | 0.946 | 1.001 | 0.276 |

We next investigated the source of this systematic undercalling of homozygous individuals across the genome. We computed the mean proportion of genotypes at every allele balance bin for common long deletions and observe an uptick of heterozygous genotypes with greater than 90% allele balance (Fig. 1a), which is markedly lower for SNVs of intermediate frequency (Fig. 1b). Overall, we find 5.4M variants at above 1% frequency where at least one individual has a heterozygous variant call with allele balance greater than 90%, and 3.2M variants (1.8M of which are indels) where this would increase the relative number of homozygous variants by more than 1%. This phenomenon is more likely to occur at common variants, where there are more opportunities for genotyping error due to the increased number of called heterozygotes; additionally, it is also more likely to occur for indels, particularly long deletions such as CCR5-Δ32 (Fig. 2a). Further, the mean increase of homozygous variant individuals using this reclassification is >5% for indels greater than 5 bp in length (Fig. 2b).

**Figure 1.**
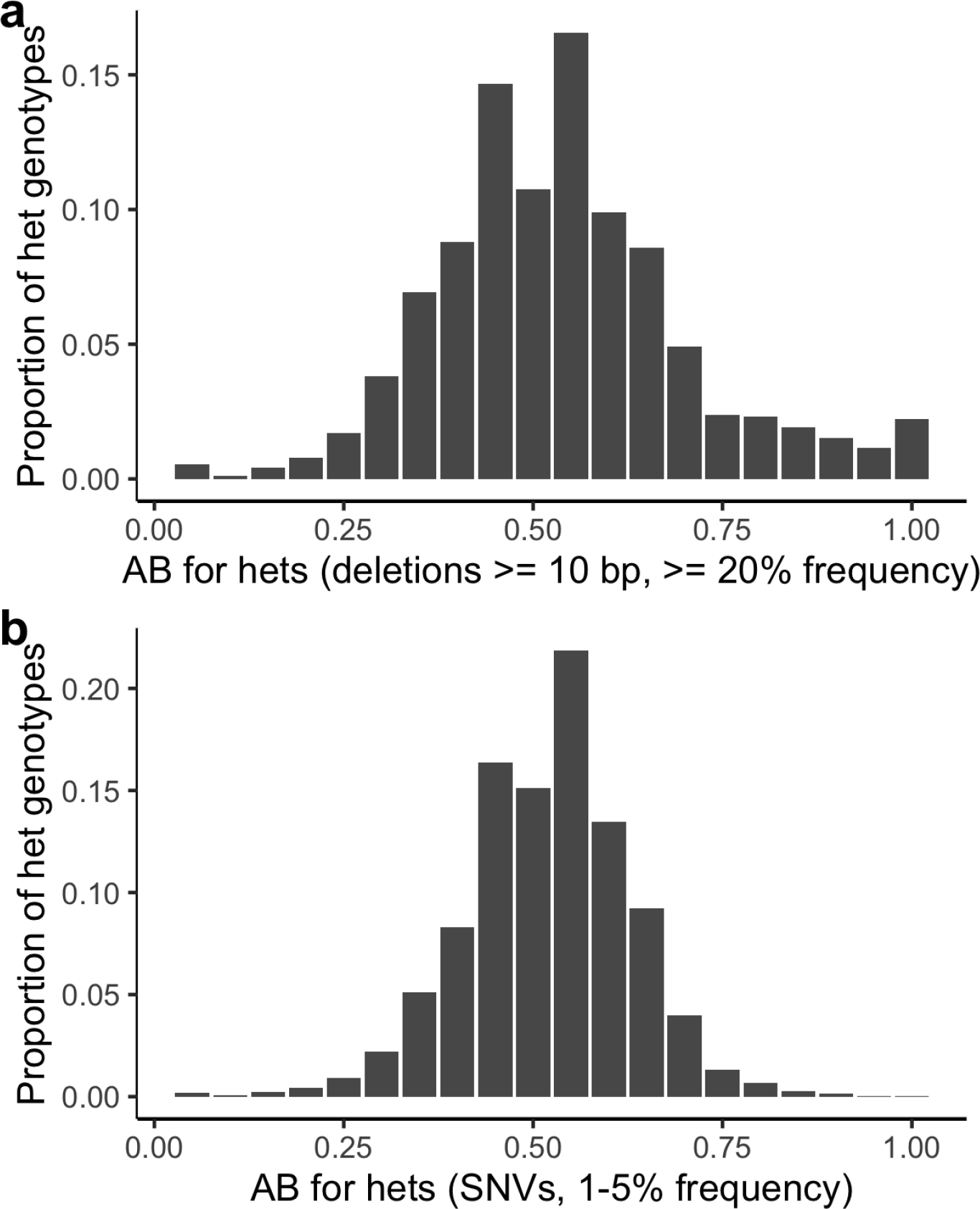
Mean proportion of genotypes at each allele balance. Previously, we computed a histogram of the allele balance for heterozygous individuals for each variant in the gnomAD genomes. Here, we show the mean of the generated histograms across all variants of a given type for common indels **(a)** and SNVs of intermediate frequency **(b)**.

**Figure 2.**
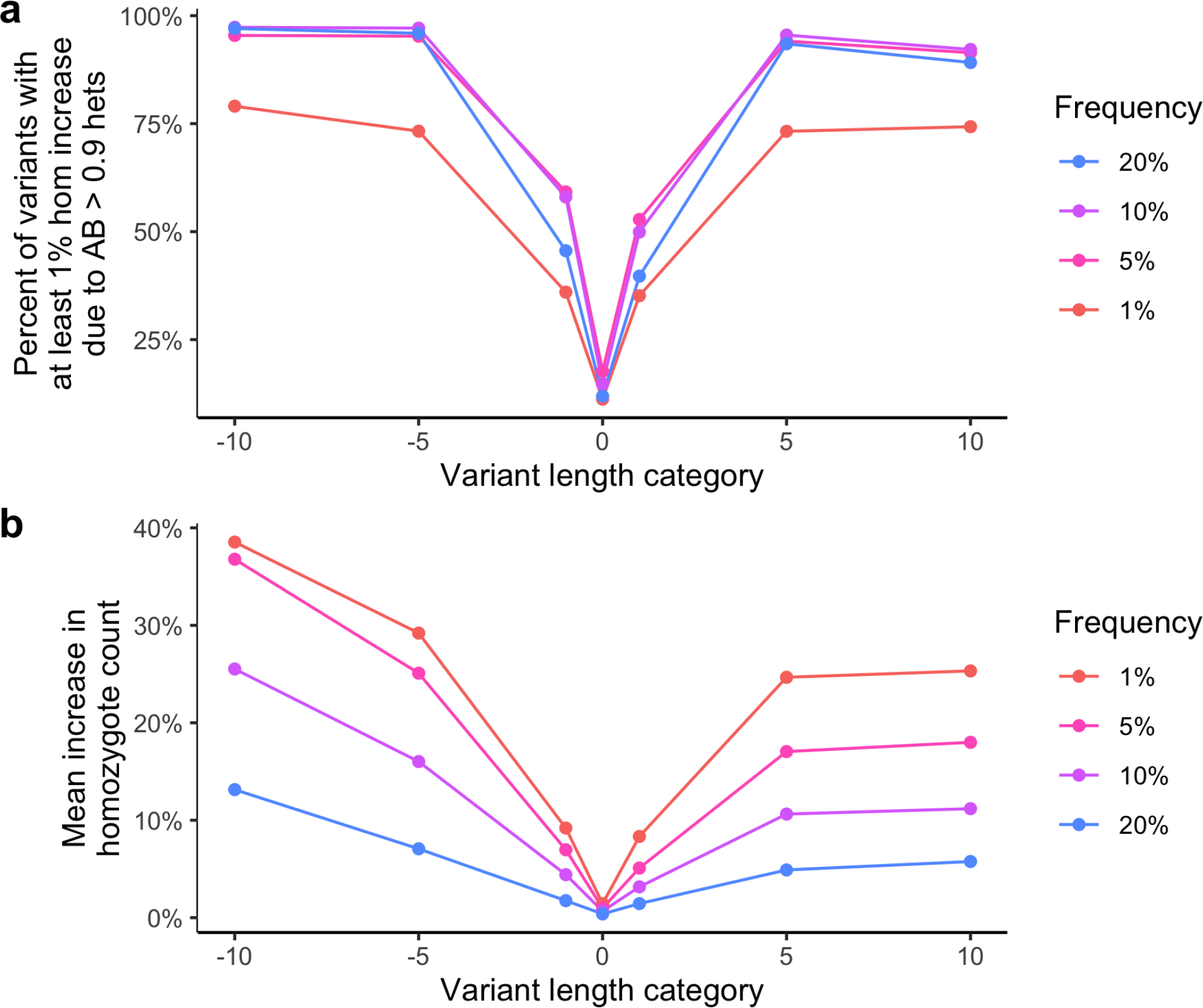
Variants and genotypes affected by high allele balance heterozygous individuals in the gnomAD genomes. **a**, The proportion of variants with at least one high allele balance heterozygous individual, split by frequency and length category. Frequency is described as the floor of the bin (i.e. 1% refers to 1-5%, 5% refers to 5-10%, 10% refers to 10-20%, 20% refers to 20-50%). Variant length category is negative for deletions (−10 refers to >= 10 bp deletions, -5 refers to 5-9 bp deletions, -1 refers to 1-4 bp deletions), 0 for SNVs, and positive for insertions (inverse of deletions). **b**, The mean percent increase in homozygous counts, split by frequency and length category. Colors and categories are consistent in **a-b**.

This artifact is a result of the likelihood model used in variant calling (see Supplement; Supplementary Fig. 1) assigning a relatively high confidence to a small number of reference-matching reads, and a lower confidence to reads that support the deletion (Supplementary Fig. 2). This artifact also occurs for SNVs at a lower rate (Fig. 2a), particularly in cases where a number of additional SNVs are found in phase, and a contaminating reference read with no mismatches provides much stronger evidence, leading to a heterozygous call (Supplementary Fig. 3).

The presence of high allele balance genotypes (0.9 < AB < 1) is consistent with sample contamination, which is common in whole exome and genome sequencing datasets. While the gnomAD dataset included a hard filter to remove samples with greater than 5% estimated contamination, lower levels of contamination may still exist and elicit these signals. Visual inspection of IGV screenshots^4^ provides evidence for this hypothesis, as in some cases, one or two high-quality reference reads are well-mapped alongside 20-30 reads with the deletion. We computed the mean number of high allele balance genotypes for each sample, and find a high correlation (r^2^ = 0.95; p < 1^-100^) between this metric and sample contamination (Fig. 3).

**Figure 3.**
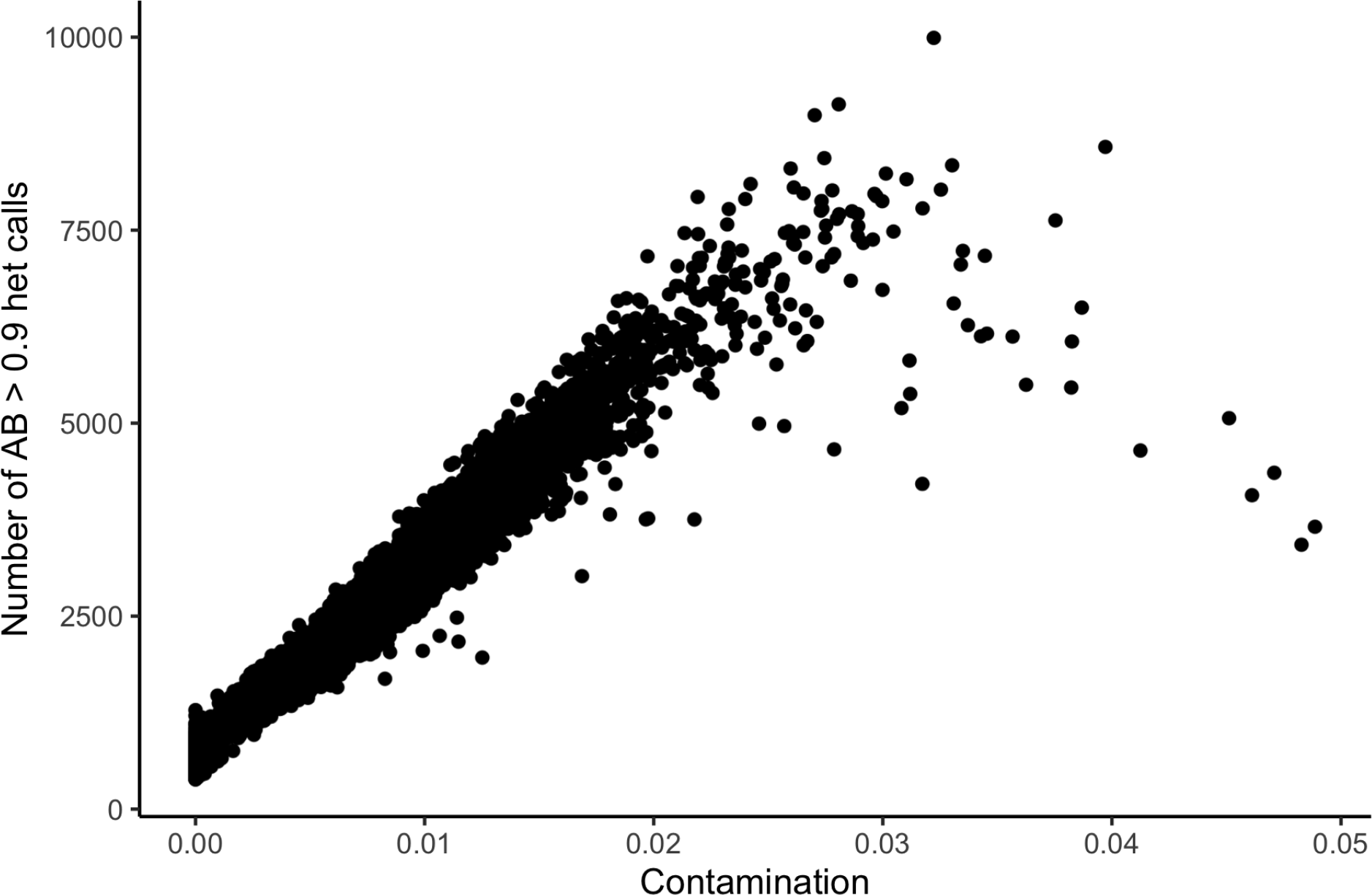
**The correlation between contamination (FREEMIX parameter from VerifyBamID^5^) and the number of high allele balance heterozygous calls in each individual in the gnomAD genomes.**

Based on these metrics, we find no evidence of depletion of homozygous CCR5-Δ32 individuals in the gnomAD dataset. We make no claims about the broader context for which the variant was originally investigated, but present this analysis as a caveat for data analysis of gnomAD and other large-scale sequencing datasets. In particular, common variation, while relatively easy to sensitively identify, is enriched for artifacts including genotyping, mapping, and annotation errors. We are grateful for the opportunity to address this overlooked error mode, and we have added a flag to the gnomAD browser to highlight variants that have underestimated the number of homozygous individuals by more than 1%. One major limitation of the current analysis is that it was performed only within the framework of the current joint-called gnomAD dataset. Further work will be required to refine the variant calling likelihood models to improve the identification of homozygous variants and to ensure this error mode can be robustly avoided in future large-scale sequencing data sets.

## Acknowledgements

We thank Valentin Ruano-Rubio, Rasmus Nielsen, Laurent Francioli, Daniel MacArthur, and Ben Neale for helpful discussions and Nicholas Watts for rapid deployment of browser fixes. We thank Kári Stefánsson, Robert Maier, Ali Akbari, Nick Patterson, and David Reich for coordinating publication of two additional manuscripts re-examining association evidence at CCR5-Δ32. After receiving an early draft of these manuscripts, Xinzhu (April) Wei and Rasmus Nielsen made the decision to retract their paper and we thank them for their integrity and commitment to the scientific method. Finally, we thank the Genome Aggregation Database (gnomAD) and the groups that provided exome and genome variant data to this resource. A full list of contributing groups can be found at https://gnomad.broadinstitute.org/about.

## Supplement

The GATK HaplotypeCaller calculates the likelihood of each read supporting an allele using the PairHMM model from Durbin et al^6^. M represents the match state (which includes SNPs), I represents an insertion in the read, and D represents a deletion in the read (Supplementary Figure 1). The transition rate delta can be thought of as a “gap open penalty” and epsilon as a “gap extension penalty”. In this way, the likelihood of large indels being derived from the reference haplotype decreases with increasing indel length. A mismatch penalty is accounted for by the emission probability of observing a particular nucleotide base in a particular alignment state, which based on the base quality estimate (not shown).

**Supplementary Figure 1.**
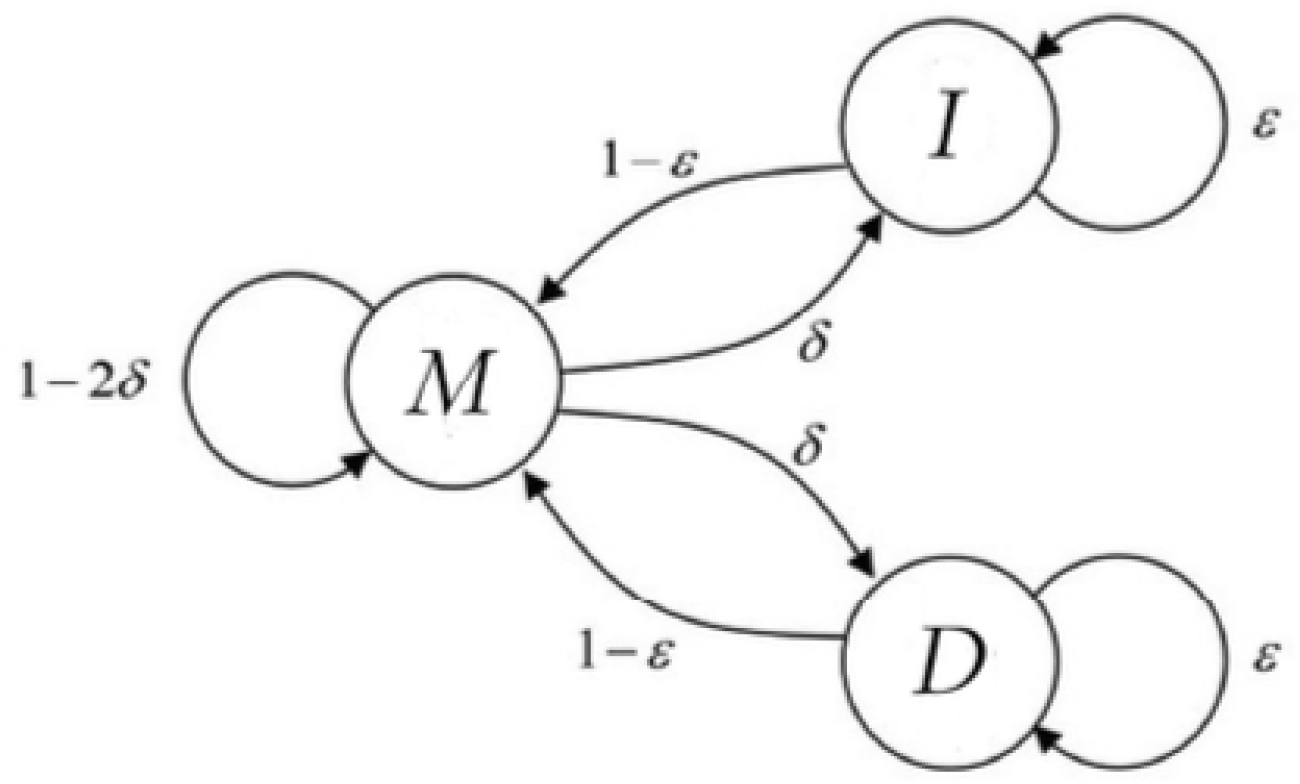
PairHMM finite state machine with transition probabilities.

Each read is assigned a likelihood based on possible alignments to each haplotype (e.g. supporting the reference or a deletion). For large indels or closely phased SNPs, a read that matches the reference is assigned a very low likelihood of being derived from the variant haplotype. However, in a non-symmetric way, the likelihood of a variant read supporting the reference is calculated as the greater of the likelihood that derives from 1) an error with alignment parameters as in the PairHMM model, or 2) a mismapping event, which is given a global probability of 10^−4.5^. This global mismapping probability limits the amount of evidence given by a variant read, whereas there is no such limitation on reference reads. At contaminated homozygous variant sites, this results in a small number of reference reads with strong support for the reference and a large proportion of alternate reads with relatively much weaker support for the variant. In this way, a small number of reference reads can result in a heterozygous genotype.

**Supplementary Figure 2.**
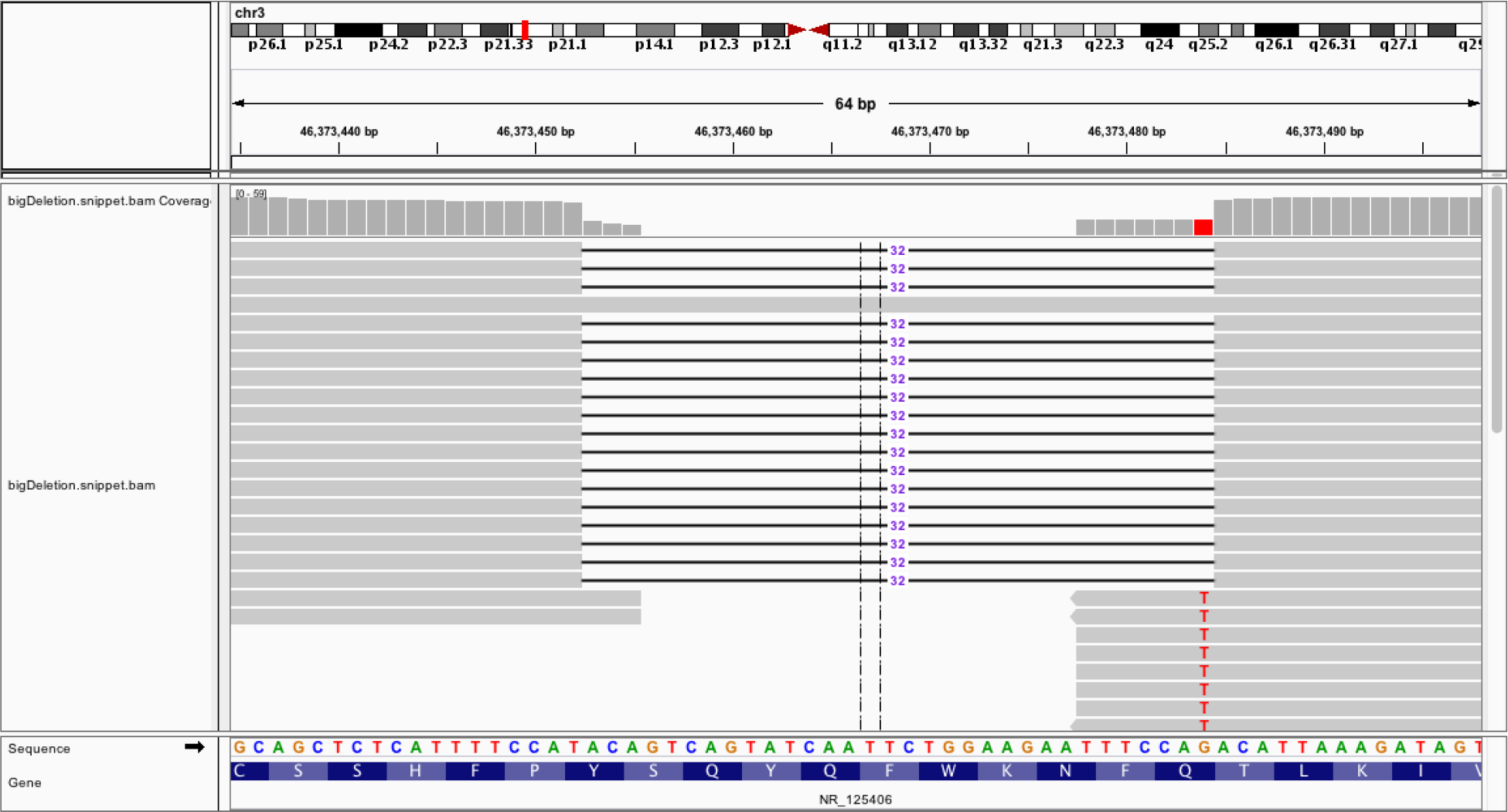
IGV visualization of reads corresponding to a called heterozygous deletion. The reported AD is 1,34,0 and PL is 1407,0,236,1410,340,1750 (both including <NON_REF> allele).

The sample shown in Supplementary Fig. 2 was sequenced with 2×151bp reads and is called as heterozygous at the position shown. The BWA-aligned pileup shows a single reference read with MQ60, CIGAR 151M. The VerifyBamID contamination estimate for this sample is 0.0155. Replacing the global mismapping probability with the contamination estimate, which is the probability of any given read being derived from another sample, and making the adjustment symmetric for both reference and alternate reads corrects the genotype to homozygous variant with PL=599,87,0,602,102,617.

We have now implemented a fix for this error mode: the maximum likelihood is now capped symmetrically for reads supporting the reference allele and reads supporting the alternate allele, preventing the types of high allele balance heterozygote calls we saw at some large deletions. The fix was released in GATK version 4.1.4.1.

**Supplementary Figure 3.**
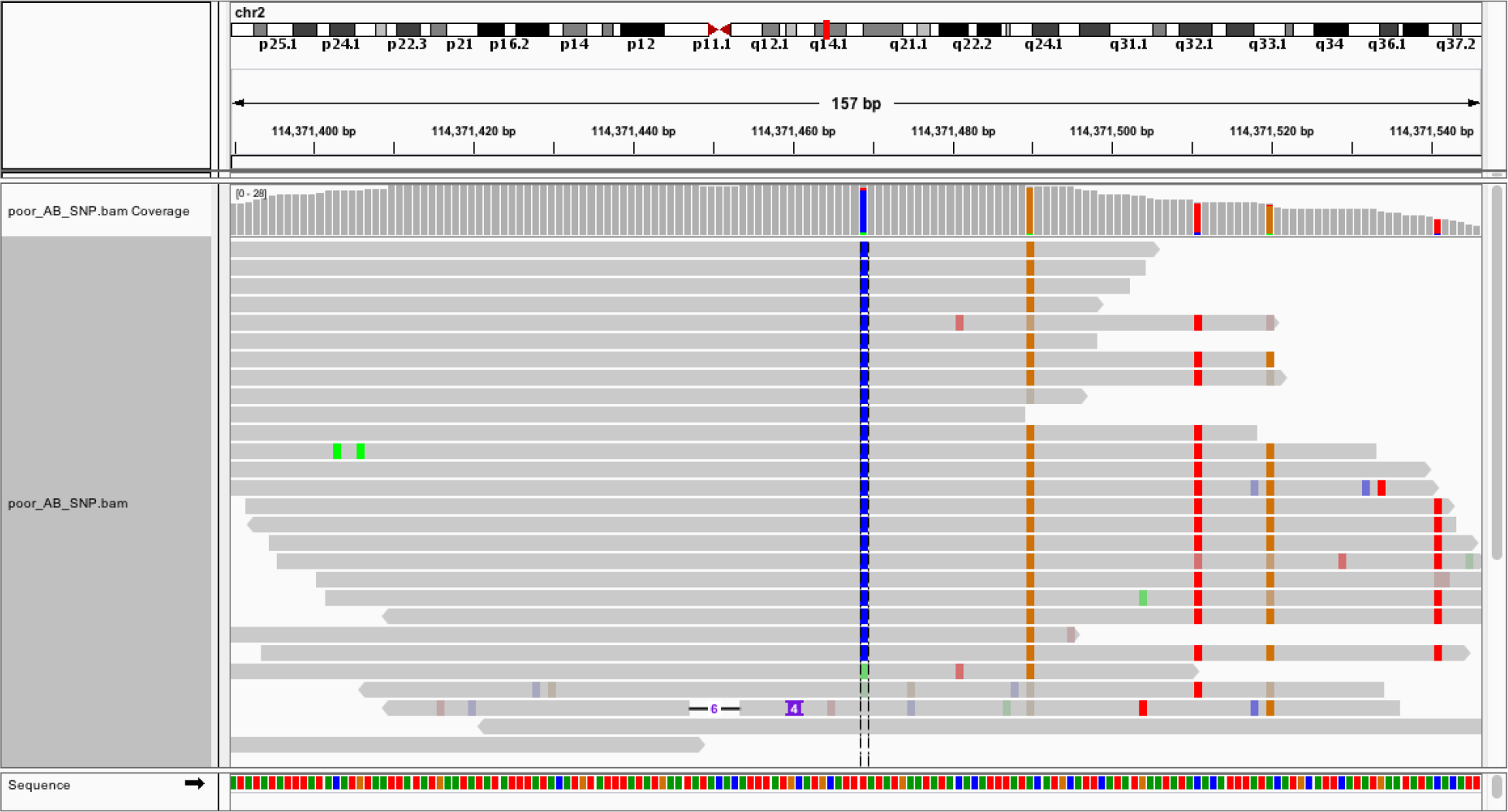
IGV visualization of reads corresponding to a called heterozygous variant. While heterozygous SNP calls with >90% variant reads are rarer, they can be explained by a similar mechanism. These SNPs occur in regions with dense phased variants such that many reads have multiple mismatches compared with the reference. The likelihood of a read derived from the reference having multiple mismatches quickly becomes smaller than the 10^−4.5^ global mismapping probability, making such SNPs subject to the same asymmetric likelihood capping artifact described for indels above.

## Notes

https://github.com/konradjk/gnomad_high_ad_hets

### Competing Interest Statement

The authors have declared no competing interest.

### Summary of Updates

Added note about implementation of fix for identified error mode

## References

1. Wei, X. & Nielsen, R. CCR5-∆32 is deleterious in the homozygous state in humans. Nat. Med. 25, 909–910 (2019).

2. Wei, X. & Nielsen, R. Deviations from Hardy Weinberg Equilibrium at CCR5-Δ32 in Large Sequencing Data Sets. bioRxiv 768390 (2019). doi:10.1101/768390

3. Karczewski, K. J., Francioli, L. C., Tiao, G. & Cummings, B. B. Variation across 141,456 human exomes and genomes reveals the spectrum of loss-of-function intolerance across human protein-coding genes. BioRxiv (2019).

4. Robinson, J. T. et al. Integrative genomics viewer. Nat. Biotechnol. 29, 24–26 (2011).

5. Jun, G. et al. Detecting and estimating contamination of human DNA samples in sequencing and array-based genotype data. Am. J. Hum. Genet. 91, 839–848 (2012).

6. Durbin, R., Eddy, S. R., Krogh, A. & Mitchison, G. Biological sequence analysis: probabilistic models of proteins and nucleic acids. (1998).

